## Supplementary Information for "Genome-wide base editor screen identifies regulators of protein abundance in yeast"

### Contents

- Supplementary Figures (Figures S1-S8) Page 2
- Supplementary Tables (Tables S7, S8, and S9) Page 10
- Supplementary Notes (Notes S1 and S2) Page 16
- Supplementary References Page 18

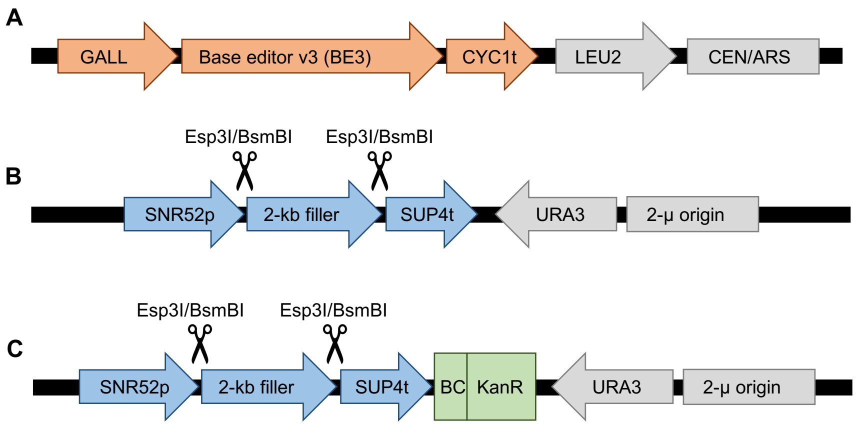

**D**

| | |
|---|---|
| gRNA plasmid backbone with 2-kb placeholder | …GCAGTGAAAGATAAATGATCGAGACG…CGTCTCTGTTTTAGAGCTAGAAATAGC…<br>…CGTCACTTTCTATTTACTAGCTCTGC…GCAGAGACAAAATCTCGATCTTTATCG… |
| Backbone digested with Esp3I/BsmBI | …GCAGTGAAAGATAAA GTTTTAGAGCTAGAAATAGC…<br>…CGTCACTTTCTATTTACTA ATCTCGATCTTTATCG… |
| Alternative 1: gRNA fragment for Golden Gate Assembly | TGATCNNNNNNNNNNNNNNNNNNNN<br>GNNNNNNNNNNNNNNNNNNNNCAAA |
| Alternative 2: gRNA fragment for Gibson Assembly | …GCAGTGAAAGATAAATGATCNNNNNNNNNNNNNNNNNNNNGTTTTAGAGCTAGAAATAGC…<br>…CGTCACTTTCTATTTACTAGNNNNNNNNNNNNNNNNNNNNCAAAATCTCGATCTTTATCG… |
| Final sgRNA plasmid | …GCAGTGAAAGATAAATGATCNNNNNNNNNNNNNNNNNNNNGTTTTAGAGCTAGAAATAGC…<br>…CGTCACTTTCTATTTACTAGNNNNNNNNNNNNNNNNNNNNCAAAATCTCGATCTTTATCG… |

**Figure S1** | Experimental resources for CRISPR base editing in *S. cerevisiae*. (**A**) The low-copy plasmid pGal-BE3 encodes the base editor v3 (BE3)[1] under a galactose-inducible promoter (GALL). This plasmid also encodes LEU2 for selection. (**B**) The high-copy pgRNA-backbone plasmid encodes a 2-kb placeholder flanked by the SNR52 promoter (a Polymerase-III promoter) and SUP4 terminator. The placeholder can be removed by Esp3I/BsmBI restriction digest and then a gRNA can be cloned in either by Golden Gate Assembly or by Gibson Assembly. This plasmid also encodes the URA3 gene for selection. (**C**) This plasmid is identical to the one shown in (B) but with a barcode cassette added. The barcode cassette contains a Kanamycin resistance gene. (**D**) Sequences at the gRNA cloning site of the pgRNA-backbone plasmid to facilitate gRNA plasmid cloning (individually by Golden Gate Assembly or as a library by Gibson Assembly). Rows 3 and 4 show the two types of DNA fragments containing the gRNA that can be cloned into the digested backbone plasmid.

**A**

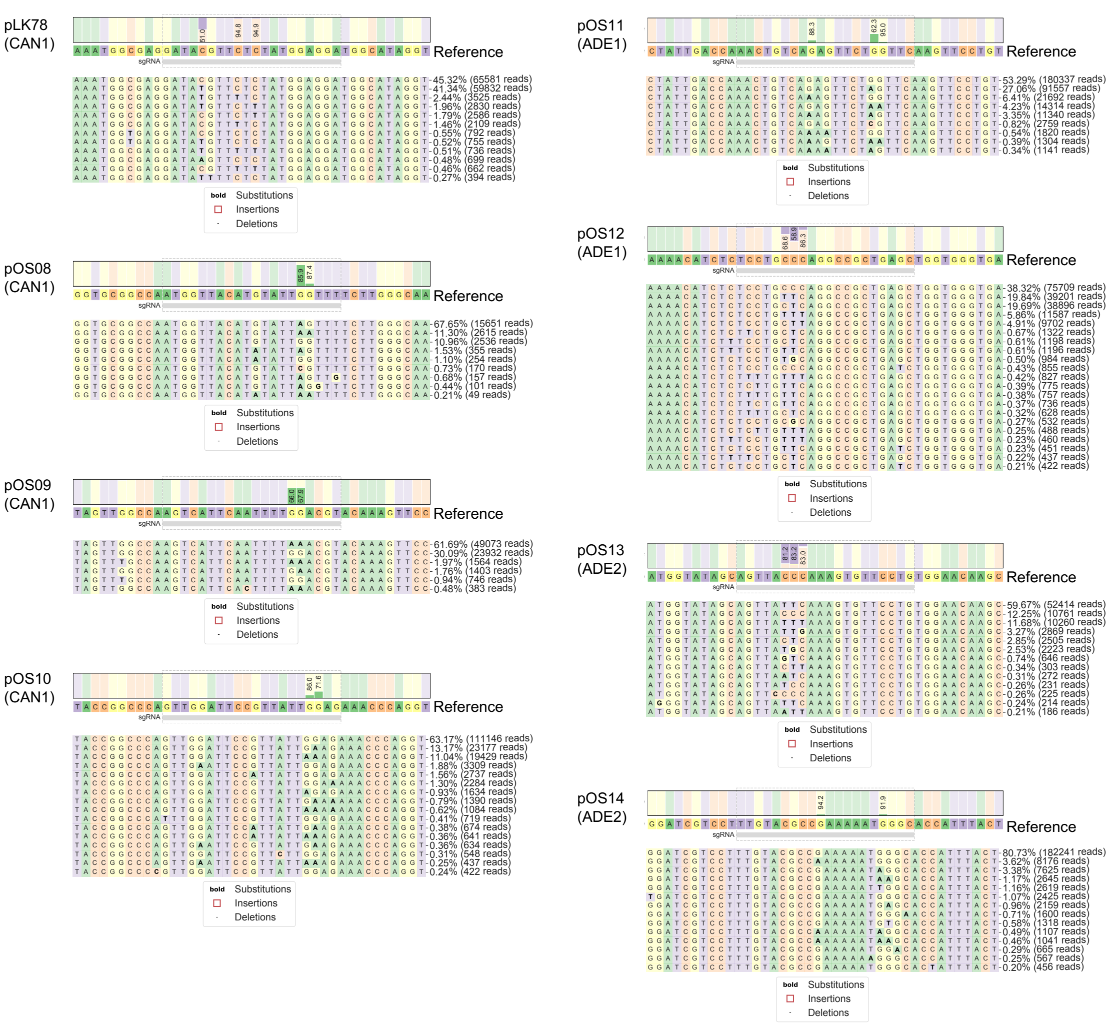

**B**

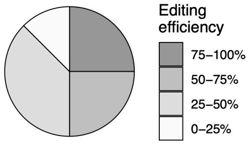

| Gene | gRNA | gRNA sequence | Editing efficiency |
|---|---|---|---|
| CAN1 | pLK78 | GATACGTTCTCTATGGAGGA | 49% |
| CAN1 | pOS08 | AAACCAATACATGTAACCAT | 86% |
| CAN1 | pOS09 | ACGTCCAAAATTGAATGACT | 68% |
| CAN1 | pOS10 | CTCCAATAACGGAATCCAAC | 28% |
| ADE1 | pOS11 | GAACCAGAACTCTGACAGTT | 38% |
| ADE1 | pOS12 | TCCTGCCCAGGCCGCTGAGC | 59% |
| ADE2 | pOS13 | AGTTACCCAAAGTGTTCCTG | 83% |
| ADE2 | pOS14 | GCCCATTTTTCGGCGTACAA | 8% |

**C**

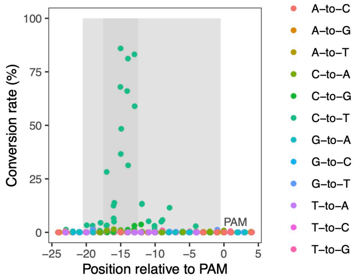

**Figure S2 |** Base editor characterization in yeast by genomic target site sequencing. Eight cultures of yeast cells each transformed with the base editor plasmid (where the base editor is under a galactose-inducible promoter) and a different gRNA plasmid were grown for 44 hours in the presence of galactose. Subsequently, the genomic target site of the gRNA in each culture was PCR-amplified and subjected to deep sequencing. (**A**) Base editing profile at the genomic target site for each of the eight gRNAs. (**B**) Classification of the eight gRNAs by editing efficiency, which reflects the maximal fraction of edits over all positions of each amplicon. (**C**) Editing outcome for the same eight gRNAs with target genomic regions aligned by PAM site. Light gray shading delineates the target site, and dark gray shading delineates the window of maximal editing.

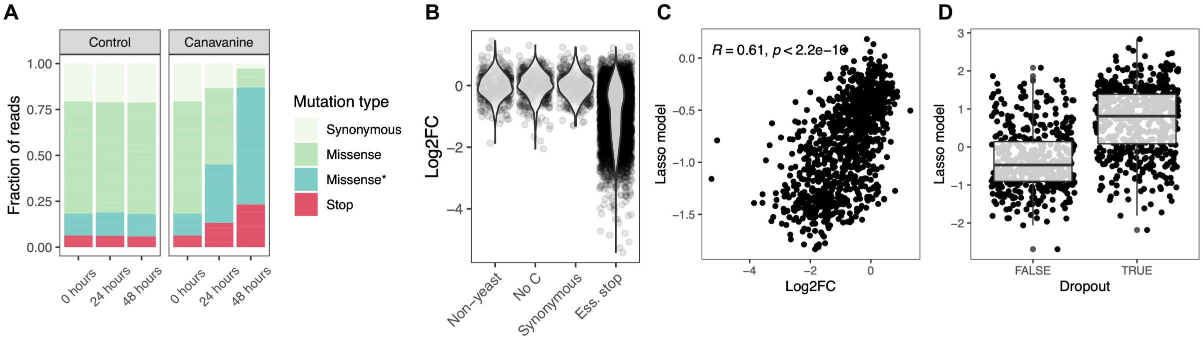

**Figure S3 |** Base editing in yeast in pooled screen format. (**A**) Fraction of reads that map to gRNAs introducing synonymous mutations, missense mutations or stop codons into the CAN1 gene in a time course experiment in which base edited yeast cultures were subjected to toxic canavanine (loss of CAN1 function renders cells resistant to canavanine) for 48 hours. The reads for the missense-introducing gRNAs are split into gRNAs without effect and gRNAs with effect (*), i.e. those that are more than two-fold enriched at 48 hours compared to 0 hours. (**B**) $Log_2$ fold change of gRNA plasmids 48 hours compared to 0 hours after induction of base editing. The gRNAs belong to either of four classes: (i) not targeting the yeast genome, (ii) no cytosine residue in the base editing window, (iii) introducing synonymous mutations only, (iv) introducing stop codons into essential genes. (**C**) Effect of sequence context on editing efficiency was assessed by applying an ordinary lasso regression model on 8000 sequence features for each of the gRNAs from classes (ii) and (iv) described in (**B**). (**D**) Same as in (**C**) but applying a logistic lasso regression model where dropout is defined as a log2FC of < -0.5.

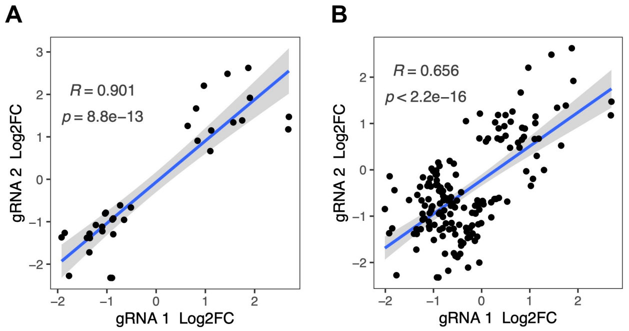

**Figure S4 |** Comparison of gRNA pairs that are predicted to target the same mutation into the genome. (**A**) All gRNA pairs for which both gRNAs have a significant (FDR < 0.05) effect on at least one protein. There are 20 gRNA pairs fulfilling this condition, some of which affect multiple proteins significantly, resulting in 33 data points. (**B**) All gRNA pairs for which at least one gRNA has a significant (FDR < 0.05) effect on at least one protein. There are 78 gRNA pairs fulfilling this condition, some of which affect multiple proteins significantly, resulting in 164 data points. Pearson correlation coefficients and corresponding p-values are indicated in the top left corner of each figure. The blue line represents the linear regression model with standard error indicated as gray shade.

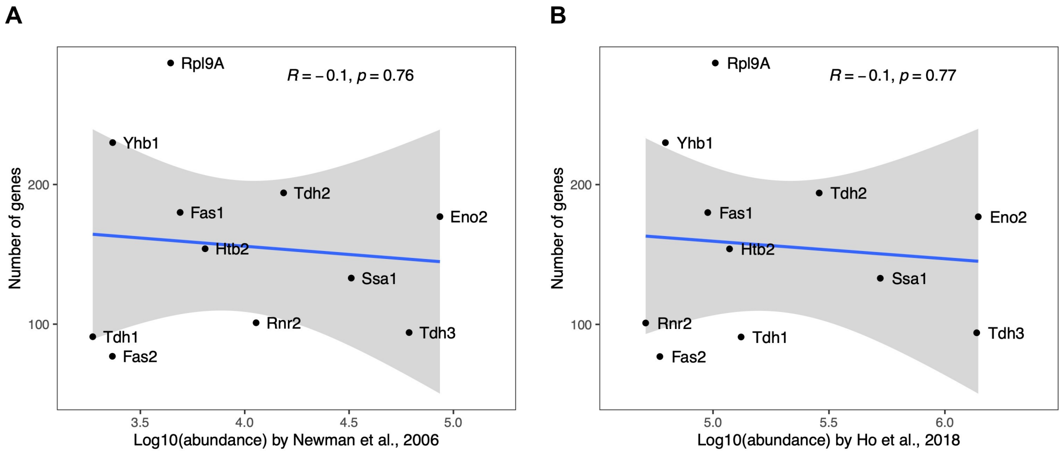

**Figure S5 |** The number of gene perturbations significantly affecting a protein (FDR < 0.05) is not dependent on the absolute abundance of that protein. Absolute protein abundances were obtained from (**A**) Newman and colleagues[2] and (**B**) Ho and colleagues[3]. Pearson correlation coefficients and corresponding p-values are indicated in the top right corner of each figure. The blue line represents the linear regression model with standard error indicated as gray shade.

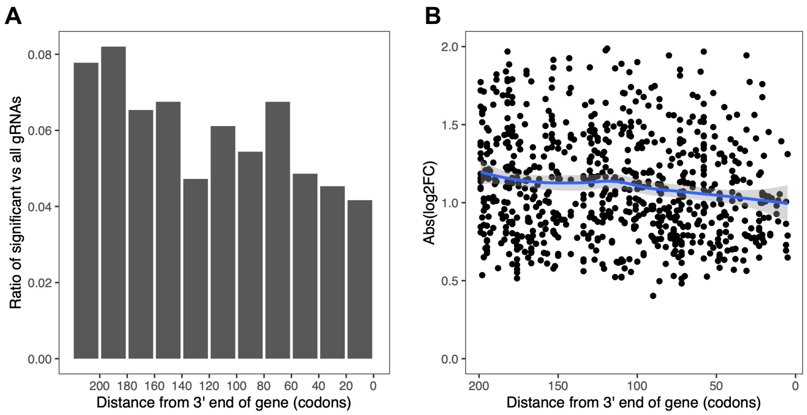

**Figure S6 |** Genetic perturbations towards the end of a gene tend to have fewer and lower effects. (**A**) Ratio of gRNAs with significant effect vs all gRNAs as a function of target gene position. (**B**) Absolute effect size of gRNAs with significant effect as a function of target gene position. The blue line shows a local polynomial regression fit (loess) with gray shades indicating the standard error. The figure was generated using the geom_smooth(method = “loess”) function from the ggplot2 R package with default parameters.

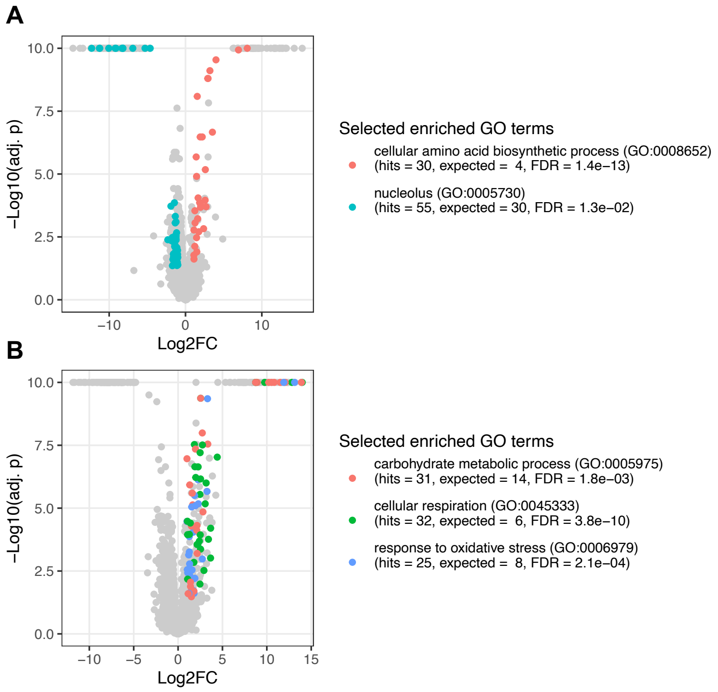

**Figure S7 |** Proteome changes in response to genetic perturbations of (**A**) POP1 (H642Y) and (**B**) SIT4 (Q184*) compared to wildtype. Selected enriched functional categories (gene ontology (GO) terms) are colored.

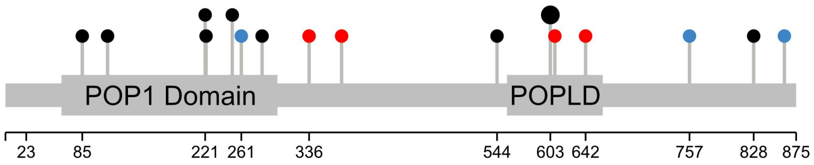

**Figure S8 |** POP1, a candidate gene for a previously identified pQTL hotspot on chromosome 14. All 14 gRNAs targeting POP1 are shown as pins along the gene. Annotated protein domains (PFAM) are indicated by wider bars. In red are gRNAs with significant effects on the abundance of at least one of the eleven proteins of interest. In blue are coding variants between the yeast strains BY S288C and RM11-1a. The larger pin head indicates two gRNAs targeting the same nucleotides. The plot was generated with the Lollipops software[4].

**Table S7** | Candidates for previously described eQTL[5] and pQTL[6] hotspots identified in segregants of a cross between the *S. cerevisiae* strains BY and RM. The candidate genes each affect at least three of our eleven proteins and contain at least one variant between BY and RM with a predicted medium or strong effect (i.e., missense mutation, indel, or premature stop codon).

| Type | Hotspot ID | Chromo some | Hotspot peak | Hotspot start | Hotspot end | Known causal gene | Candidate genes (this study) |
|---|---|---|---|---|---|---|---|
| pQTL | 2 | chrII | | 89002 | 227097 | | PRE7; RIB1; PET9 |
| pQTL | 3 | chrII | | 369902 | 415106 | | SEC18; UBC4 |
| pQTL | 7 | chrVII | | 92196 | 186694 | | ROK1; SUA5 |
| pQTL | 10 | chrVIII | | 413697 | 452100 | | PRP8; DBP8 |
| pQTL | 11 | chrX | | 107906 | 178103 | | SSY5 |
| pQTL | 14 | chrXII | | 125601 | 297999 | | DRS1; LMO1; PPR1 |
| pQTL | 18 | chrXIV | | 200799 | 293198 | | JJJ1; POP1; RAP1; YNL198C; CHS1 |
| pQTL | 19 | chrXIV | | 415298 | 530699 | MKT1 | DBP2; ARP5 |
| pQTL | 20 | chrV | | 106200 | 203100 | IRA2 | WRS1; ADH1; IRA2; THP1 |
| eQTL | 3 | chrI | 146738 | 138499 | 167326 | | SEN34 |
| eQTL | 5 | chrII | 13061 | 13061 | 18004 | | PKC1 |
| eQTL | 15 | chrIII | 262264 | 255891 | 264282 | | SRB8 |
| eQTL | 34 | chrVI | 156375 | 132559 | 161152 | | SPB4 |
| eQTL | 35 | chrVI | 229348 | 227693 | 240289 | | SAP155 |
| eQTL | 38 | chrVII | 194505 | 174975 | 205155 | | ROK1; SUA5; PMR1 |
| eQTL | 40 | chrVII | 397449 | 397449 | 398572 | OLE1 | OLE1 |
| eQTL | 43 | chrVII | 851062 | 850923 | 876915 | | TYS1 |
| eQTL | 55 | chrX | 187765 | 181214 | 191872 | | SPT10; GCD14 |
| eQTL | 57 | chrX | 429703 | 416411 | 455247 | | CTK2; CYR1; SUI2 |
| eQTL | 62 | chrXI | 102908 | 102671 | 109811 | | FAS1; PRS1 |
| eQTL | 71 | chrXII | 856042 | 845725 | 873750 | | ADE13; SSQ1 |
| eQTL | 81 | chrXIV | 18838 | 18046 | 24221 | | EGT2 |
| eQTL | 86 | chrXV | 50257 | 41495 | 62744 | | RIB4; RTC1 |
| eQTL | 88 | chrXV | 172956 | 168708 | 181326 | IRA2 | IRA2 |
| eQTL | 98 | chrXVI | 207364 | 207364 | 209279 | | PPQ1 |

### **Table S8** | Yeast Strains

| Name | Mutation | Background | Notes |
|---|---|---|---|
| yOS057, yOS058, yOS059 | RAS2 S46L | S288C BY4741 (MATa his3Δ1 leu2Δ0 met15Δ0 ura3Δ0) | |
| yOS061, yOS062, yOS063 | RAS2 Q272* | S288C BY4741 (MATa his3Δ1 leu2Δ0 met15Δ0 ura3Δ0) | |
| yOS054, yOS055, yOS056 | IRA2 W342* | S288C BY4741 (MATa his3Δ1 leu2Δ0 met15Δ0 ura3Δ0) | |
| yOS079, yOS080, yOS082 | SSY5 A637K/R | S288C BY4741 (MATa his3Δ1 leu2Δ0 met15Δ0 ura3Δ0) | |
| yOS064, yOS065, yOS066 | POP1 H642Y | S288C BY4741 (MATa his3Δ1 leu2Δ0 met15Δ0 ura3Δ0) | |
| yOS074, yOS075, yOS076 | SIT4 Q184* | S288C BY4741 (MATa his3Δ1 leu2Δ0 met15Δ0 ura3Δ0) | |
| GFP strains | GOI-GFP(S65T)-HIS3MX6 | S288C BY4741 (MATa his3Δ1 leu2Δ0 met15Δ0 ura3Δ0) | Gene of interest (GOI) is C-terminally tagged with a GFP(S65T) cassette containing also HIS3 from *L. kluyveri* [12] |

**Table S9** | Plasmids

| Name | Main plasmid component | gRNA targeting sequence | Note |
|---|---|---|---|
| pGal-BE3 (original name: pOS02) | Base editor | n/a | Plasmid deposited with Addgene (##172409) |
| pgRNA-backbone (original name: pOS05a) | gRNA cassette | 2-kb placeholder from lentiCRISPR v2 plasmid obtained from Addgene #52961 [68] | Plasmid deposited with Addgene (#172517) |
| pLK78 | CAN1 T64M gRNA | GATACGTTCTCTATGGAGGA | Plasmid obtained from Addgene (#43803) [65] |
| pOS08 | CAN1 W174* gRNA | AAACCAATACATGTAACCAT | |
| pOS09 | CAN1 W195* gRNA | ACGTCCAAAATTGAATGACT | |
| pOS10 | CAN1 W263* gRNA | CTCCAATAACGGAATCCAAC | |
| pOS11 | ADE1 W64* gRNA | GAACCAGAACTCTGACAGTT | |
| pOS12 | ADE1 Q178* gRNA | TCCTGCCCAGGCCGCTGAGC | |
| pOS13 | ADE2 Q122* gRNA | AGTTACCCAAAGTGTTCCTG | |
| pOS14 | ADE2 W185* gRNA | GCCCATTTTTCGGCGTACAA | |
| pOS34 | RAS2 S46L gRNA | GGATTCATACAGGAAGCAAG | |
| pOS35 | RAS2 Q272* gRNA | CAACGCTCAAAGCGCTAATA | |
| pOS29 | IRA2 W342* gRNA | TATCCAAAACATTATTGCTT | |
| pOS40 | SSY5 A637T gRNA | TCCCCCGCACTAGCAAATAA | |
| pOS32 | POP1 H642Y gRNA | CATCCATAACACTAAATTAC | |
| pOS39 | SIT4 Q184* gRNA | GCTCAAGAAGTGCCACACGA | |

### **Table S10** | Primers

| Name | Description | Sequence | Note |
| --- | --- | --- | --- |
| OS123 | Barcoding KanR cassette fwd | GTGGTGCTTTTTTTGTTTTTTATGTCTNNNNNNNNNNNNNNNNNNNNCCGGAATTGCCAGCTGGG | 20-nucleotide barcode |
| OS124 | Barcoding KanR cassette rev | GCGCGTAATACGACTCACTATAGGCTGTATGCGGTGTGAAATACC | |
| OS088 | Backbone for BC-KanR cassette fwd | CCCTATAGTGAGTCGTATTACGCGC | |
| OS089 | Backbone for BC-KanR cassette rev | AGACATAAAAAACAAAAAAAGCACCAC | |
| OS113 | Sublibrary CanAde fwd | TGGGTGATTCCGCAGGGTCCAGAGT | |
| OS116 | Sublibrary CanAde rev | ACTGGTCAATGCGAAGGAAACGCGT | |
| OS111 | Sublibrary ES fwd | TGGGTGATTCCGCGTCGAGTAGGGT | |
| OS112 | Sublibrary ES rev | ACTGGTCAATGCCGTGTGAAGCTGG | |
| OS152 | Sublibrary EP fwd | TGGGTGATTCGGTCGAGCCGGAACT | |
| OS153 | Sublibrary EP rev | ACTGGTCAATGGATGCGCACCCAGA | |
| OS150 | Sublibrary NS fwd | TGGGTGATTCCGATCGCCCTTGGTG | |
| OS151 | Sublibrary NS rev | ACTGGTCAATGGTTTAGCCGGCGTG | |
| OS154 | Sublibrary NP fwd | TGGGTGATTCTCCCGGCGTTGTCCT | |
| OS155 | Sublibrary NP rev | ACTGGTCAATGCTCCGTCACTGCCC | |
| OS119 | Sublibrary 2nd PCR fwd | TGAAAGTTGGTGCGCATGTTTCGGCGTTCGAAACTTCTCCGCAGTGAAAGATAAATGATC | |
| OS120 | Sublibrary 2nd PCR rev | ACTTTTTCAAGTTGATAACGGACTAGCCTTATTTTAACTTGCTATTTCTAGCTCTAAAAC | |
| OS125 | gRNA-barcode sequencing rev | TCGTCGGCAGCGTCAGATGTGTATAAGAGACAGtaagtagagGCGTTCGAAACTTCTCCGCAG | PAGE-purified |
| OS130 | gRNA-barcode sequencing 1 fwd | TCGTCGGCAGCGTCAGATGTGTATAAGAGACAGtaagtagagGGCTAGCGGTAAAGGTGCGC | Used individually, not as a pool<br><br>Note that these primers resulted in unexpected PCR products and were later replaced by OS140-149. |
| OS131 | gRNA-barcode sequencing 2 fwd | TCGTCGGCAGCGTCAGATGTGTATAAGAGACAGatcatgcttaGGCTAGCGGTAAAGGTGCGC | |
| OS132 | gRNA-barcode sequencing 3 fwd | TCGTCGGCAGCGTCAGATGTGTATAAGAGACAGgatgcacatctGGCTAGCGGTAAAGGTGCGC | |
| OS133 | gRNA-barcode sequencing 4 fwd | TCGTCGGCAGCGTCAGATGTGTATAAGAGACAGcgattgctcgacGGCTAGCGGTAAAGGTGCGC | |
| OS134 | gRNA-barcode sequencing 5 fwd | TCGTCGGCAGCGTCAGATGTGTATAAGAGACAGtcgatagcaattcGGCTAGCGGTAAAGGTGCGC | |
| OS135 | gRNA-barcode sequencing 6 fwd | TCGTCGGCAGCGTCAGATGTGTATAAGAGACAGatcgatagttgcttGGCTAGCGGTAAAGGTGCGC | |

| | | | |
|---|---|---|---|
| OS136 | gRNA-barcode sequencing 7 fwd | TCGTCGGCAGCGTCAGATGTGTATAAGAGACAGgatcgatccagttagGGCTAGCGGTAAAGGTGCGC | |
| OS137 | gRNA-barcode sequencing 8 fwd | TCGTCGGCAGCGTCAGATGTGTATAAGAGACAGcgatcgatttgagcctGGCTAGCGGTAAAGGTGCGC | |
| OS138 | gRNA-barcode sequencing 9 fwd | TCGTCGGCAGCGTCAGATGTGTATAAGAGACAGacgatcgatacacgatcGGCTAGCGGTAAAGGTGCGC | |
| OS139 | gRNA-barcode sequencing 10 fwd | TCGTCGGCAGCGTCAGATGTGTATAAGAGACAGtacgatcgatggtccagaGGCTAGCGGTAAAGGTGCGC | |
| OS140 | gRNA-barcode sequencing 1 fwd | TCGTCGGCAGCGTCAGATGTGTATAAGAGACAGtaagtagagGCGTTCGAAACTTCTCCGCAG | Used as pool |
| OS141 | gRNA-barcode sequencing 2 fwd | TCGTCGGCAGCGTCAGATGTGTATAAGAGACAGatcatgcttaGCGTTCGAAACTTCTCCGCAG | |
| OS142 | gRNA-barcode sequencing 3 fwd | TCGTCGGCAGCGTCAGATGTGTATAAGAGACAGgatgcacatctGCGTTCGAAACTTCTCCGCAG | |
| OS143 | gRNA-barcode sequencing 4 fwd | TCGTCGGCAGCGTCAGATGTGTATAAGAGACAGcgattgctcgacGCGTTCGAAACTTCTCCGCAG | |
| OS144 | gRNA-barcode sequencing 5 fwd | TCGTCGGCAGCGTCAGATGTGTATAAGAGACAGtcgatagcaattcGCGTTCGAAACTTCTCCGCAG | |
| OS145 | gRNA-barcode sequencing 6 fwd | TCGTCGGCAGCGTCAGATGTGTATAAGAGACAGatcgatagttgcttGCGTTCGAAACTTCTCCGCAG | |
| OS146 | gRNA-barcode sequencing 7 fwd | TCGTCGGCAGCGTCAGATGTGTATAAGAGACAGgatcgatccagttagGCGTTCGAAACTTCTCCGCAG | |
| OS147 | gRNA-barcode sequencing 8 fwd | TCGTCGGCAGCGTCAGATGTGTATAAGAGACAGcgatcgatttgagcctGCGTTCGAAACTTCTCCGCAG | |
| OS148 | gRNA-barcode sequencing 9 fwd | TCGTCGGCAGCGTCAGATGTGTATAAGAGACAGacgatcgatacacgatcGCGTTCGAAACTTCTCCGCAG | |
| OS149 | gRNA-barcode sequencing 10 fwd | TCGTCGGCAGCGTCAGATGTGTATAAGAGACAGtacgatcgatggtccagaGCGTTCGAAACTTCTCCGCAG | |
| LG_CAN1_1F | Genomic region CAN1 T64M fwd | TCGTCGGCAGCGTCAGATGTGTATAAGAGACAGAGACGCCGACATAGAGGAGA | |
| LG_CAN1_1R | Genomic region CAN1 T64M rev | GTCTCGTGGGCTCGGAGATGTGTATAAGAGACAGACCAAGGGCAATCATACCAA | |
| LG_CAN1_2F | Genomic region CAN1 W174* fwd | TCGTCGGCAGCGTCAGATGTGTATAAGAGACAGTTGGTATGATTGCCCTTGGT | |
| LG_CAN1_2R | Genomic region CAN1 W174* rev | GTCTCGTGGGCTCGGAGATGTGTATAAGAGACAGAAGTTCCAGGGCAAAAGTGA | |
| LG_CAN1_3F | Genomic region CAN1 W195* fwd | TCGTCGGCAGCGTCAGATGTGTATAAGAGAC | |

| | | |
|---|---|---|
| | | AGTCACTTTTGCCCTGGAACTT |
| LG_CAN1_3R | Genomic region CAN1 W195* rev | GTCTCGTGGGCTCGGAGATGTGTATAAGAGACAGGCACCTGGGTTTCTCCAATA |
| LG_CAN1_4F | Genomic region CAN1 W263* fwd | TCGTCGGCAGCGTCAGATGTGTATAAGAGACAGGTTTGTGGTGCTGGGGTTAC |
| LG_CAN1_4R | Genomic region CAN1 W263* rev | GTCTCGTGGGCTCGGAGATGTGTATAAGAGACAGTCTTGGAACGGATTTTCTGG |
| OS070 | Genomic region ADE1 W64* fwd | TCGTCGGCAGCGTCAGATGTGTATAAGAGACAGTGCTGTTTGTTGCTACGGATCG |
| OS071 | Genomic region ADE1 W64* rev | GTCTCGTGGGCTCGGAGATGTGTATAAGAGACAGAGAGAGCGGTCTTCTAGTTGCG |
| OS072 | Genomic region ADE1 Q178* fwd | TCGTCGGCAGCGTCAGATGTGTATAAGAGACAGTCCCAGAACCAATCTTCACCCC |
| OS073 | Genomic region ADE1 Q178* rev | GTCTCGTGGGCTCGGAGATGTGTATAAGAGACAGTAGAGGAGTCTGGCGTTAGCAC |
| OS074 | Genomic region ADE2 Q122* fwd | TCGTCGGCAGCGTCAGATGTGTATAAGAGACAGCCAATGACCACGTTAATGGCTCC |
| OS075 | Genomic region ADE2 Q122* rev | GTCTCGTGGGCTCGGAGATGTGTATAAGAGACAGTCAATAGGGACGTCTCACTGGC |
| OS076 | Genomic region ADE2 Q185* fwd | TCGTCGGCAGCGTCAGATGTGTATAAGAGACAGCGAGGACTTTGGCATACGATGG |
| OS077 | Genomic region ADE2 Q185* rev | GTCTCGTGGGCTCGGAGATGTGTATAAGAGACAGCGCCTTAAGTTGAACGGAGTCC |
| OS242 | Genomic region RAS2 S46L fwd | GTACTTGCCGCCTTCGTCTCTA |
| OS243 | Genomic region RAS2 S46L rev | TGGTGGTGGTGGTGTTGGTAAA |
| OS244 | Genomic region RAS2 Q272* fwd | TCTTGGAGGCTTCACTGGTGTT |
| OS245 | Genomic region RAS2 Q272* rev | GTCGTGAATGCCAGGAATGCAA |
| OS232 | Genomic region IRA2 W342* fwd | CAAGTGCCCGTAACACAGACAC |
| OS233 | Genomic region IRA2 W342* rev | CGAAGGGGAAGATGAGCCTTGA |
| OS252 | Genomic region SSY5 A637T fwd | GCAACATACCAACAAGCCCCAA |
| OS253 | Genomic region SSY5 A637T rev | GGGCGAGAGAGCAATCGTAGAT |
| OS238 | Genomic region POP1 H642Y fwd | TCGACAACGACATAGCCTTGGT |
| OS239 | Genomic region POP1 H642Y rev | GTGTTACCATGGTTCTGGCTGC |
| OS250 | Genomic region SIT4 Q184* fwd | CTGACGCTGGCCGCTATAATTG |
| OS251 | Genomic region SIT4 Q184* rev | TCGACCTTCATTACACTCGCGA |

### Note S1 | Base editing in yeast is efficient and predictable

To adapt the base editor for use in yeast, we created a plasmid which contains the base editor under a galactose-inducible promoter, as well as a plasmid optimized for bulk cloning of gRNA libraries (Figure S1). We first assessed base-editing efficiency and target window in yeast by amplicon-sequencing the genomic target loci of eight individual gRNAs. We observed the expected C-to-T mutations in over 50% of the reads for four of the eight gRNAs tested, and in over 25% of the reads for three additional gRNAs (Figure S2). Furthermore, we found that the target window and mutagenesis pattern are very similar to those described in human cells—95% of edits are C-to-T transitions, and 89% of these occurred in a five-nucleotide region 13 to 17 base pairs upstream of the PAM sequence (Figure S2)[1].

To assess base editor performance in a pooled screen format, we used a library of all 90 gRNAs suitable for targeting the CAN1 gene (Table S1). Loss-of-function mutations in CAN1 render yeast resistant to the toxic arginine analog canavanine. We exposed base-edited yeast cultures to canavanine for 48 hours and observed a strong enrichment of reads from gRNAs predicted to introduce premature stop codons (chi-squared test, $\chi^2$ = 1737, df = 1, p < 2.2e-16), as well as a depletion of reads from gRNAs predicted to introduce synonymous mutations in CAN1 (chi-squared test, $\chi^2$ = 1637, df = 1, p < 2.2e-16) (Figure S3A, Table S2). Eleven of the 65 gRNAs predicted to introduce missense mutations resulted in an enrichment similar to that of the stop codons. The amino acid substitutions introduced by these 11 gRNAs therefore likely disrupt the function of CAN1. This inference is supported by their higher conservation as reflected by more negative PROVEAN scores[7]; 7 of the 11 amino acid substitutions have a PROVEAN score less than -5, vs. 11 of 54 for gRNAs not showing enrichment (Fisher's exact test, OR = 0.15, p = 0.007).

To further characterize base editing efficiency in a pooled screen format, we designed a library containing all 5430 gRNAs predicted to introduce stop codons into essential genes, together with three sets of 500 control gRNAs each (random gRNA sequences not targeting the yeast genome, gRNAs without a C in the targeting region, gRNAs introducing synonymous mutations) (Table S1). We evaluated the effects of these gRNAs on yeast survival over 48 hours (Table S3). As expected, cells transformed with control gRNAs remained in the culture at stable levels, whereas cells transformed with gRNAs introducing stop codons into essential genes tended to drop out over time (59% displayed $\log_2$ fold change < -0.5 and false-discovery rate (FDR) < 0.05) (Figure S3B). We asked whether DNA sequence context in the targeted region could explain which gRNAs introduced stop codons with high efficiency. We built a lasso regression model with over 8000 sequence features per gRNA and found that it could explain 37% of the observed variance in gRNA depletion (Pearson's R = 0.61, p < 2.2e-16) (Figure S3C and D). In particular, the base preceding the target C has a strong effect—a T or another C produces much higher editing rates than a G. These findings in yeast are in line with recent reports in human cell lines and can help guide the design of future gRNA libraries[8,9].

**Note S2 | Base editor screen identifies causal gene candidates underlying regulatory hotspots in natural yeast strains**

To date, the most comprehensive picture of genetic architecture underlying gene expression regulation in yeast has been achieved by linkage mapping in crosses of divergent yeast strains[5]. A remarkable finding is that a majority of genetic loci affecting mRNA and protein expression (eQTLs and pQTLs, respectively) cluster in regulatory hotspots. However, these regulatory hotspots can be large genomic regions encompassing up to dozens of genes each, making it challenging to pinpoint the causal gene and, hence, to understand their mechanism of action. With the base editor screen, we were able to identify for 24 of these regulatory hotspots a total of 47 candidate causal genes, each affecting three or more of the eleven proteins and harboring at least one parental variant with a predicted medium to strong effect (Table S7). For only three of these 24 regulatory hotspots the causal gene is known and our screen identified two of them, IRA2 and OLE1, as candidate causal genes[10,11]. Yet unknown causal gene candidates include POP1, which overlaps a protein regulatory hotspot on chromosome 14 and harbors several coding variants in natural yeast strains (Figure S8)[6]. Future studies using denser tiling of gRNAs in CRISPR-based screens or deep mutational scanning will reveal the role of each residue for POP1 function and confirm whether POP1 is indeed the causal gene underlying this regulatory hotspot in natural yeast strains. Overall, this example showcases how targeted genetic screens, especially with methods that give nucleotide-level resolution like the CRISPR base editor, can provide insights into how natural genetic variants affect gene and protein regulation.
